# Lower shear stress exacerbates atherosclerosis by inducing the generation of neutrophil extracellular traps *via* Piezo1-mediated mechanosensation

**DOI:** 10.1101/2023.02.19.529165

**Authors:** Ying Zhu, Tian Wang, Zining Wang, Xiaohui Chen, Liu Wang, Ruyan Niu, Zixin Sun, Chong Zhang, Yang Luo, Yijie Hu, Wei Gu

## Abstract

**BACKGROUND:** Atherosclerosis is a chronic lipid-driven inflammatory disease, largely influenced by hemodynamics. Neutrophil extracellular traps (NETs)-mediated inflammation plays an important role in atherosclerosis. However, little is known about the mechanism of the generation of NETs under different shear stress and subsequent damage to endothelial cells. We sought to identify a novel mechanical signal provokes NETs generation and to investigate its potential role in atherosclerosis.

**METHODS:** ApoE^−/−^ mice were fed with high-fat diet (HFD) to induce atherosclerosis. The model of lower shear stress (LSS) with a partial ligation of the left carotid artery was established to assess the role of LSS in NETs generation and atherosclerotic lesions development. Furthermore, the underlying mechanism of LSS promoting NETs generation and injuring endothelial cells was deciphered in neutrophil-like human promyelocytic leukemia (HL-60) cells in parallel-plate flow chamber.

**RESULTS:** We found that LSS correlated spatially with both NETs and atherosclerosis, while inhibition of NETosis could significantly reduce plaque formation in ApoE^−/−^ mice. *In vitro*, LSS could promote NETs generation directly through down-regulation of Piezo1, a mechanosensitive ion channel. downexpression of Piezol could activate neutrophils and promote NETosis in static. Conversely, Yoda1-evoked activation of Piezo1 attenuated LSS-induced NETosis. Mechanistically, the downexpression of Piezo1 resulted in decreased Ca^2+^ influx and increased histone deacetylase 2 (HDAC2), which increase reactive oxygen species levels, then led to NETosis. LSS-induced NETs generation promoted the apoptosis and adherence of endothelial cells.

**CONCLUSIONS:** LSS directly promotes NETosis through piezo1-HDAC2 axis in atherosclerosis progression. This study uncovers the essential role of Piezo1-mediated mechanical signaling in NETs generation and plaque formation, which provides a promising therapeutic strategy for atherosclerosis.

**Graphic Abstract:** 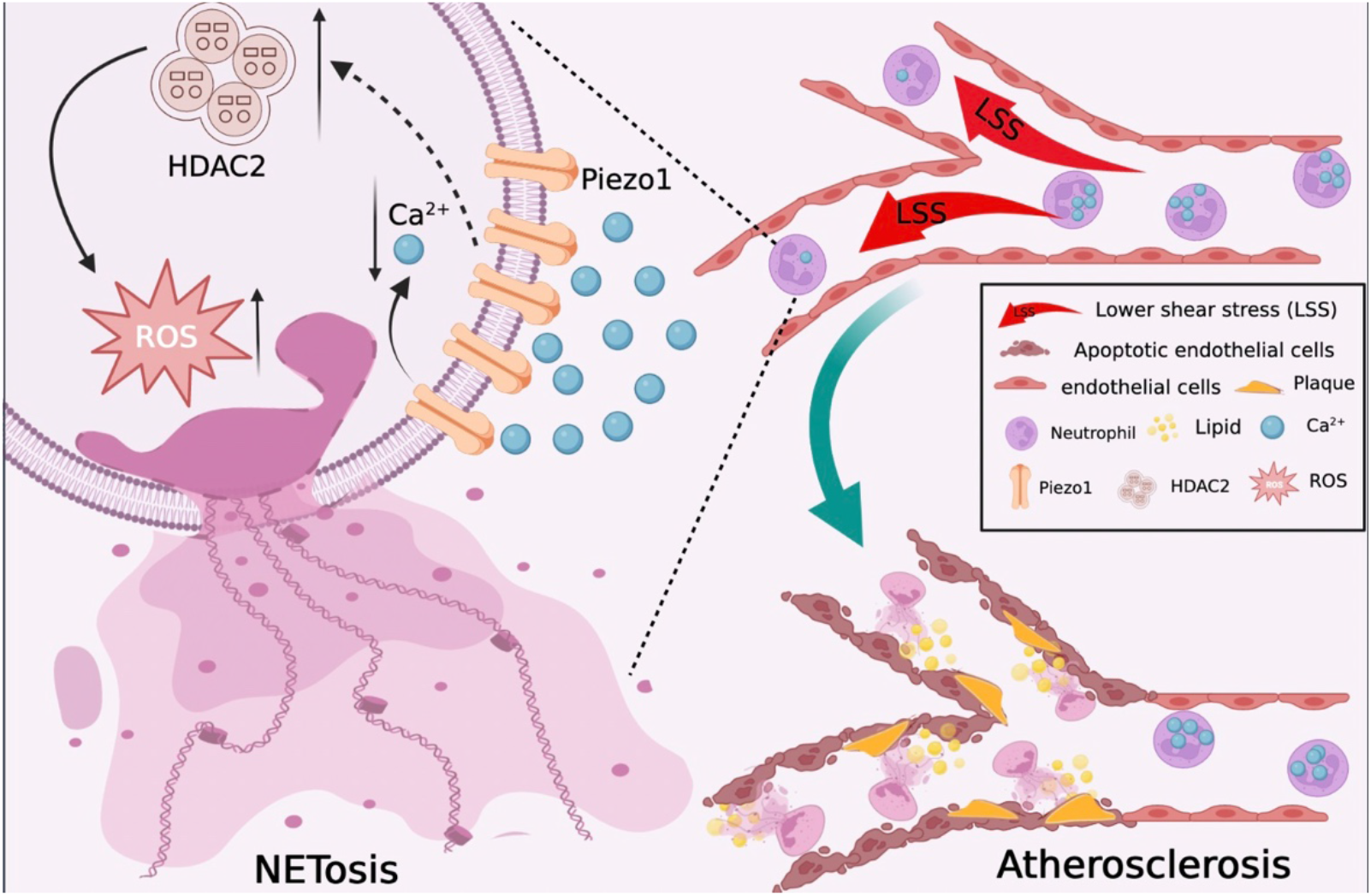

Proposed mechanism for lower shear stress LSS exacerbating atherosclerosis. LSS stimuli decrease Piezo1 expression in the neutrophils, resulting in decreased intracellular Ca^2+^ concentration, as well as the higher expression level of HDAC2, which could activate oxidative stress and promote intracellular reactive oxygen species formation, and ultimately lead to NETs generation. NETs could aggravate endothelial cells injury and exasperate atherosclerosis.

**Highlights:**

■ Lower shear stress (LSS) promotes Neutrophil extracellular traps (NETs) formation, which is critical for lipid deposits and plaque formation in Atherosclerosis.
■ Atherosclerotic plaque formation was significantly reduced in the aorta of high fat diet fed ApoE^−/−^ mice intraperitoneal injected with NETs inhibitor, GSK484, especially in the lower shear stress regions.
■ Piezo1 is a key molecule in the process of neutrophils sense lower shear stress.
■ lower shear stress inhibits the activation of Piezo1 and promotes NETosis through piezo1-HDAC2 axis.
■ LSS-induced NETs promote the apoptosis and adhesion of endothelial cells.

## 1. Introduction

Atherosclerosis is the most common pathological basis of cardiovascular disease and leads to mortality and morbidity worldwide^1, 2^. Many life-threatening cardiovascular events, such as thrombosis, ischaemic heart disease, and stroke are mainly caused by the rupture of atherosclerotic plaques inside the arterial wall. However, there is no conclusion as to what triggers the formation of atherosclerotic plaques.

Multiple factors have been found to participate in atherosclerosis, including endothelial dysfunction, oxidized low-density lipoproteins (oxLDLs), inflammation, and oxidative stress. Recent evidence support that neutrophils, the most-abundant innate immune sentinels, play an important role in atherosclerosis^3^. One of the possible mechanisms is related to NETosis, In this process, activated neutrophils release neutrophil extracellular traps (NETs) which are fiber-like structures composed of decondensed chromatin and protein granules. It has been found that ApoE^−/−^ mice injected with lipopolysaccharide (LPS) produced larger lesion sizes with the accumulation of myeloid cells and NETs^4^. NETs are also found to be highly accumulated in the downstream plaque regions with vulnerability features^5^. Patients with unstable plaques have increased levels of peptidyl arginine deiminase-4 (PAD4) compared to patients with stable plaques^6^. Either a PAD4 deficiency in bone marrow-derived cells or administration of DNase I could significantly alleviate arterial intimal injury and plaque formation^7^, suggesting that NETs are involved in the initial stage of atherosclerosis. However, the triggers of NETosis in atherosclerosis are still under investigation.

Previous studies have shown that various stimuli can activate neutrophils, which is crucial for NETosis. NETs are released by neutrophils exposed to gram positive and negative bacteria, in particular large pathogens and fungi^8^. The virulence factors of bacteria prime NETs release *via* secretion of interleukin-6 (IL-6), tumor necrosis factor-α (TNF-α), and reactive oxygen species (ROS)^9^, or forming pores on the neutrophils’ nuclear membrane and inducing the release of DNA bearing histones and granular proteins including myeloperoxidase (MPO) and neutrophil elastase (NE) into the extracellular space^10, 11^. Activated platelets can also induce NETs formation *via* P-selectin (CD62P) and P-selectin glycoprotein ligand 1 (PSGL-1)^12^. In addition, NETs formation is triggered by other distinct stimuli including nitric oxide, urate crystals, autoantibodies, proinflammatory cytokines, and the interaction of neutrophils with damaged endothelial cells^13^.

Meanwhile, atherosclerosis is a typical hemodynamic-related disease. The lesions prefer to localize within characteristic vessel geometries, such as the inside of the vessel curvature, downstream of stenosis, on the sidewall of a vessel bifurcation, and adjacent to the opening of a branch, which is the regions of low shear stress (LSS) and disturbed flow^14^. In addition, LSS impairs endothelial cells’ function through a variety of signaling pathways^15, 16^. The biomechanical damage effect of blood flow shear stress has attracted increasing attention^17^. However, whether LSS can directly trigger NETosis is still unknown.

In the study we assessed the role of LSS in NETs generation and atherosclerotic lesions development *in vivo*, and investigated the underlying mechanism of LSS promoting NETs generation and injuring endothelial cells in neutrophil-like HL-60 (dHL-60) cells in parallel-plate flow chamber. The results found that the LSS-related NETs formation promoted atherosclerosis and might be a promising therapeutic target in the future.

## 2. Materials and Methods

The Supplemental Methods and the Major Resources Table can be found in the Supplemental Material.

### 2.1 Animals

All animal experiments were performed under specific pathogen-free barrier conditions following institutional guidelines, and the protocol was approved by the Committee on the Ethics of Animal Experiments of Chongqing University (Permit Number: CQU-IACUC-RE-202210-001). ApoE^−/−^ mice (4-6 weeks old, male) were purchased from Cavenslasales Company (Changzhou, China). They were kept in a pathogen-free animal care facility under the standard laboratory conditions (26 ± 2 ^°^C, 40%-60% humidity, 12-h light/dark cycle) and allowed full access to fresh water and food. Mice were randomly distributed into four groups (n = 5/group): (i) normal chow diet (NCD) for 4 weeks, (ii) NCD for 8 weeks, (iii) HFD for 4 weeks, and (iv) HFD for 8 weeks. To further study the causal role of NETs on plaque formation, HFD-fed mice (n = 5/group) were intraperitoneally injected with PAD4 inhibitors, 10 mg/kg GSK484 (#HY-100514, MedChem Express, New Jersey, USA), or vehicle control (saline) daily for 8 weeks. To analyze neutrophil activation by lower flow *in vivo*, ApoE^−/−^ mice underwent partial ligation of the left carotid artery (LCA) as previously described^18^.

Blood was collected by cardiac puncture of deeply anesthetized mice. Plasma and serum were separated by centrifugation (2,000×g for 10 min at 4 ^°^C) respectively and stored at -80 ^°^C before use. Triglycerides (TG), total cholesterol (TC), high-density lipoprotein cholesterol (HDL-C), and low-density lipoprotein cholesterol (LDL-C) levels in serum were measured using a Chemray 240 automatic chemistry analyzer (Rayto, Shenzhen, China). The IL-6 levels in plasma were analyzed using a Mouse IL-6 Uncoated ELISA kit (#88-7064, Invitrogen, MA, USA) according to the manufacturer’s instructions. Neutrophil count was measured using an automatic animal blood cell analyzer (#BC-2800vet, Mindray, Shenzhen, China). For organ harvest mice were sacrificed by cervical dislocation. Hearts, aortas, and carotid arteries were then dissected and perfusion fixed with 4% paraformaldehyde for further analysis.

### 2.2 Cell culture, differentiation and drug treatment

Human promyelocytic leukemia (HL-60) cells and human umbilical vein endothelial cells (HUVECs) were grown in RPMI-1640 medium (Gibco, Thermo Scientific, MA, United States) supplemented with 10% fetal bovine serum (FBS) (Gibco, Thermo Scientific, MA, United States), and 1% antibiotics (100 IU/ml penicillin, 100 mg/ml streptomycin), in a humidified atmosphere containing 5% CO_2_ at 37 ^°^C. The dHL-60 cells were prepared by treating HL-60 cells with 1.3% dimethyl sulfoxide (DMSO, #D3650, Sigma-Aldrich, MO, United States) for 5 days. Neutrophil-like phenotype was assessed by Wright-Giemsa stain (#C0135, #C0133, Beyotime Biotechnology, Shanghai, China) and flow cytometry based on CD11b staining (#101205, Biolegend, CA, USA). The dHL-60 cells were incubated with the Piezo1 agonist Yoda1 (1 μM, #HY-18723, MedChem Express, New Jersey, USA) for 24 h, the HDAC2 inhibitor valproic acid sodium (VPA, 5 mM, #HY-10585A, MedChem Express, New Jersey, USA), or agonist Theophylline (5 μM, #HY-B0809, MedChem Express, New Jersey, USA) for 6h, respectively.

### 2.3 Parallel-plate flow chamber assay

Parallel-plate flow chamber assays were performed with dHL-60 cells as described before^19^. Briefly, 15-20 ml suspensions with 10^6^-10^7^ dHL-60 cells were transferred in a sterile glass bottle and perfused with the flow by a peristaltic pump, The shear stress was controlled by the flow rate, and the formula τ= 6Qμ/wh^2^ was used to calculate the fluid shear stress (τ), Q is the flow rate and μ is the dynamic viscosity of the perfusate. dHL-60 cells were perfused at the shear rate of 5 dyne/cm^2^ (LSS), 12 dyne/cm^2^ (, physiological shear stress, PSS), or 0 dyne/cm^2^ (Static) for 24 h using a parallel plate flow chamber system.

### 2.4 Immunofluorescence staining

For immunofluorescence staining, arotid artery and proximal aortic sections and cells-plated slides were fixed with 4% paraformaldehyde for 30 min, permeabilized with 0.2% Triton X-100/PBS at 4 ^°^C for 20 min, and then blocked with 10% normal donkey serum (Jackson Immuno Research, PA, USA) for 1 hour, following by primary antibodies at 4 ^°^C overnight and secondary antibody in a blocking reagent for 2 hours at room temperature (RT). After rinsing with PBST, nuclei were co-stained with DAPI (#C1005, Beyotime Biotechnology, Shanghai, China) and slides were mounted. Microscopy was performed with Zeiss laser scanning confocal microscopes (LSM 980, Jena, Germany).

Primary antibodies: Human/Mouse Myeloperoxidase/MPO Antibody, (#AF3667), R&D Systems, MN, USA, (1:200); Anti-Histone H3 (citrulline R2 + R8 + R17) antibody, (#ab5103), Abcam, Cambridge, UK, (1:200); Purified anti-mouse Ly-6G Antibody, (#127601), Biolegend, CA, USA, (1:200); Anti-FAM38A/PIEZO1 antibody, (#ab128245), Abcam, Cambridge, UK, (1:200); HDAC2 Polyclonal antibody, (#12922-3-AP), Proteintech, Wuhan, China, (1:200); CD31 Monoclonal antibody, (#66065-2-Ig), Proteintech, Wuhan, China, (1:200).

Secondary antibodies: Donkey Anti-Rabbit IgG(H+L), FITC conjugate, (#SA00003-8), Proteintech, Wuhan, China, (1:200); Donkey Anti-Goat IgG(H+L), Cy3 conjugate, (#SA00009-3), Proteintech, Wuhan, China, (1:200); Alexa Fluor 647 AffiniPure Donkey Anti-Mouse IgG (H+L), (#34113ES60), Yeasen, Shanghai, China, (1:200).

### 2.5 Flow cytometry assays

To analyze the efficiency of dHL-60 differentiation, cell suspensions were obtained. After 500×g centrifugation for 5 min at 4 ^°^C, the pellet was resuspended in PBS and incubated for 30 min at 4 ^°^C with rat FITC-conjugated mAb against CD11b (1:200, BioLegend, CA, USA). After incubation, cells were centrifuged at 500×g (4 ^°^C for 5 min), washed with PBS and resuspended.

For intracellular Ca^2+^ detection, cell suspensions were obtained. After 500×g centrifugation for 5 min at 4 ^°^C, the pellet was resuspended in PBS and incubated for 30 min at 37 ^°^C with the Ca^2+^-sensitive indicator Fluo-4 AM (#S1060, Beyotime Biotechnology, Shanghai, China) at a final concentration of 1 μM.

ROS detection was performed using a ROS detection kit (#MA0219, Meilun, Dalian, China) followed by the manufacturer’s instructions. Briefly, cell suspensions were collected and stained with the ROS probe 2,7-dichloride-hydro fluorescein diacetate (DCFH-DA) at 37 ^°^C for 30 min. After incubation, cells were centrifuged at 500×g (4 ^°^C for 5 min), washed with PBS and resuspended.

For apoptosis detection, a total of 1×10^6^ HUVECs cells were collected after co-cultured with shear stress-induced dHL-60 cells. The early and late apoptosis (Annexin-V^+^ PI^-^ plus Annexin-V^+^ PI^+^) were measured *via* an apoptosis detection Kit (#MA0428, Meilun, Dalian, China) according to the manufacturer’s instructions.

The dHL-60 cells differentiation, intracellular Ca^2+^, ROS and cell apoptosis assay were all measured *via* flow cytometry (B90883, Backman Coulter, CA, USA), and the data were analyzed using FlowJo 10.0.

### 2.6 Statistical Analysis

All quantitative results are presented as the means ± SEM. The quantification of atherosclerotic lesions area in the aorta *En face* was performed by Image J. GraphPad Prism software (Version 8.0) was used for statistical analysis. An unpaired t-test was used for normally distributed data to compare differences between two groups, and one-way ANOVA with Bonferroni’s test was used for multiple comparisons. *P* < 0.05 was considered statistically significant.

## 3. Result

### 3.1 Lower shear stress correlates spatially with both NETs and atherosclerosis

To explore the effect of LSS on the generation of NETs and atherosclerotic lesions, we established mouse models of atherosclerosis by feeding ApoE^−/−^ mice with HFD for 4 to 8 weeks. *En face* Oil-Red O staining showed prominent lipid deposition in vessel wall at 4 weeks and more obviously at 8 weeks, especially at the area of the bifurcation (Figure. 1A and 1B). TG, TC, HDL-C, LDL-C, and IL-6 concentrations, as well as the proportion of neutrophils, were all significantly increased in HFD groups after 4 weeks (Figure. 1C). To assess the effect of LSS on neutrophils *in vivo*, we generated a partial ligation model in the left carotid artery (LCA) of ApoE^−/−^mice (Figure. 1D). This model only leaves open the occipital artery (OA) and generates lower shear stress in LCA compared with the right carotid artery (RCA). Anti-Ly-6G, MPO, and citrullination of histone H3 (CitH_3_) immunofluorescence stainings were performed in the blood vessel’s frozen sections after 2-4 weeks of treatment. In HFD mice, partial LCA ligation induced advanced atherosclerotic lesions assessed in the ligated carotid, with abundant lipid deposition. Interestingly, more NETs were found in the lipid lesions of LCA compared with RCA, indicating that the LSS had an effect on NETs generation. Additionally, the accumulation of NETs significantly increased in carotid artery with the prolongation of modeling time after 4 weeks of treatment (Figure. 1E).

**Figure 1.**
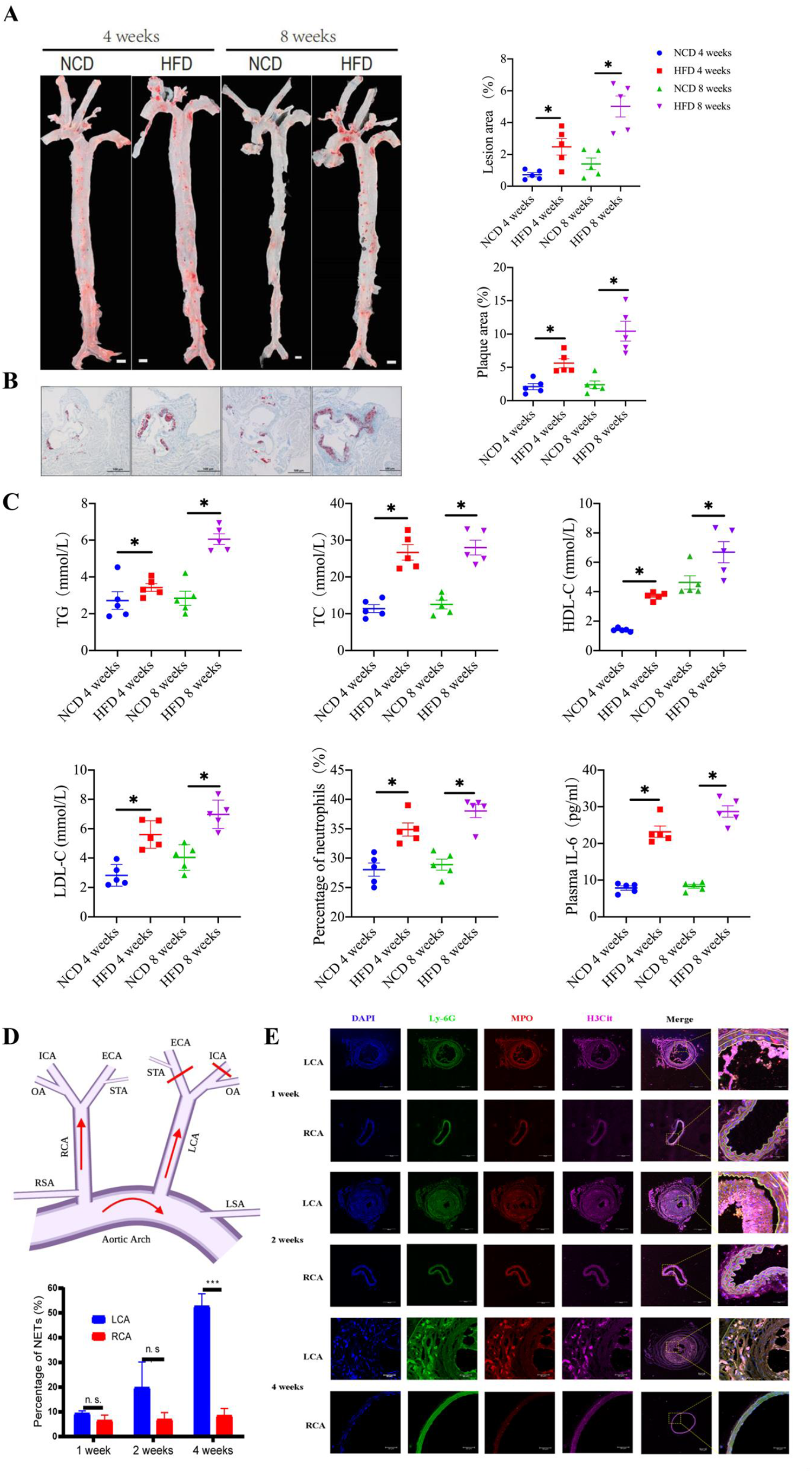
LSS promoted neutrophil extracellular traps (NETs) generation *in vivo*. (A) *En face* stained aortaventralis and quantification of lipid deposition within lesion area. Scale bar,1 mm. (B) Oil Red O stained aortic root and quantification of plaque size demonstrated as the percentage of lesion area within aortic root area. Scale bar, 300 μm. (C) The concentration of TG, TC, HDL-C, LDL-C, and IL-6 concentration and the proportion of neutrophils in whole blood. (D) The model of partial ligation. (E) Immunostain analysis of the level of NETs Quantitative analysis of the percentage of numbers of CitH3-positive cells relative to DAPIstained cells and the mean fluorescence intensity of CitH3. The data are presented as the means ±SE. (n=5). ns: *P* > 0.05, \**P* < 0.05, \*\*\**P* < 0.001.

### 3.2 Inhibition of NETosis reduces plaque formation

To demonstrate the direct relationship between NETs generation and atherosclerotic plaques formation, 4-week-old ApoE^−/−^ mice were fed with HFD and injected intraperitoneally (i.p.) with GSK484 daily to inhibit NETs formation. Daily injection of saline served as control (Figure. 2A). *En face* staining of aortaventralis showed less lipid deposition on the blood vessels of the GSK484 group compared with control group, especially at vascular bifurcations (Figure. 2B). H&E and Oil-Red O staining results both showed less progression of atheromatous plaque in GSK484 group (Figure. 2C and 2D). Immunofluorescence staining of carotid artery and proximal aortic sections showed less NETs location on the intima of the blood vessels of the GSK484 group compared with control group (Figure. S1A and S1B). Interestingly, there was no significant difference in serum TG, TC, HDL-C, and LDL-C concentrations and proportion of neutrophils between the two groups, while the GSK484 group have higher plasma IL-6 levels (Figure. 2E and 2F).

**Figure 2.**
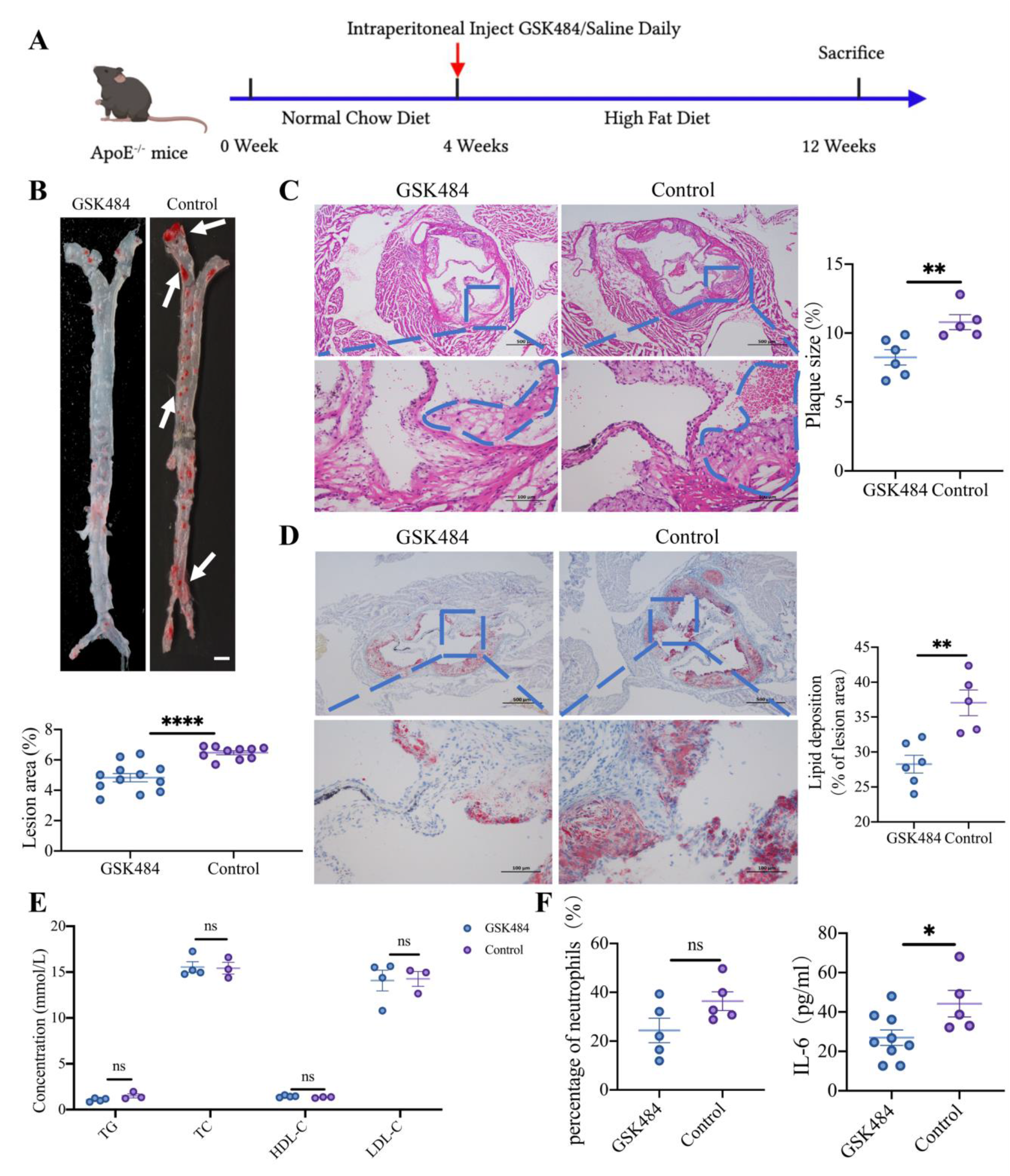
Suppression neutrophil extracellular traps (NETs) generation mitigates atherosclerotic plaques *in vivo*. ApoE*^−/−^* mice were divided into GSK484 and Control group. (A) Schematic diagrams of the drug delivery mouse model. (B) *En face* staining of aortas. Scale bar: 1mm. Plaque area as a percentage of the total area was quantified (n=10). (C and D) H&E and Oil Red O staining of aortic roots (n=5). Quantification of plaque size, Oil Red O positive area in plaque. Scale bar: 100 μm. (E) Serum TG, TC, HDL-C, LDL-C concentration. (F) Plasma IL-6 concentration and the proportion of neutrophils in whole blood. The data are presented as the mean ± SEM. ns: *P* > 0.05, \**P* < 0.05, \*\**P* < 0.01, \*\*\**P* < 0.001, \*\*\*\**P* < 0.0001.

### 3.3. LSS promotes NETs generation In Vitro and aggravates endothelial cells apoptosis

dHL-60 cells were confirmed by light microscopy of Wright-Giemsa-stained cytospin specimens and flow cytometry (Figure. S2A and S2B). We further examined the neutrophils’ responses to different shear stress in dHL-60 cells using a parallel-plate flow chamber (Figure. 3A). The number of viable cells decreased with increased treatment time, especially in the LSS group (Figure. S2C). Triple immunofluorescent staining with anti-MPO, anti-CitH3 antibodies, and DAPI confirmed that CitH3^+^ cells were significantly increased in LSS group. This result was further confirmed by Western blot analysis, implying the formation of NETs could be induced by LSS directly (Figure 3B and 3C).

**Figure 3.**
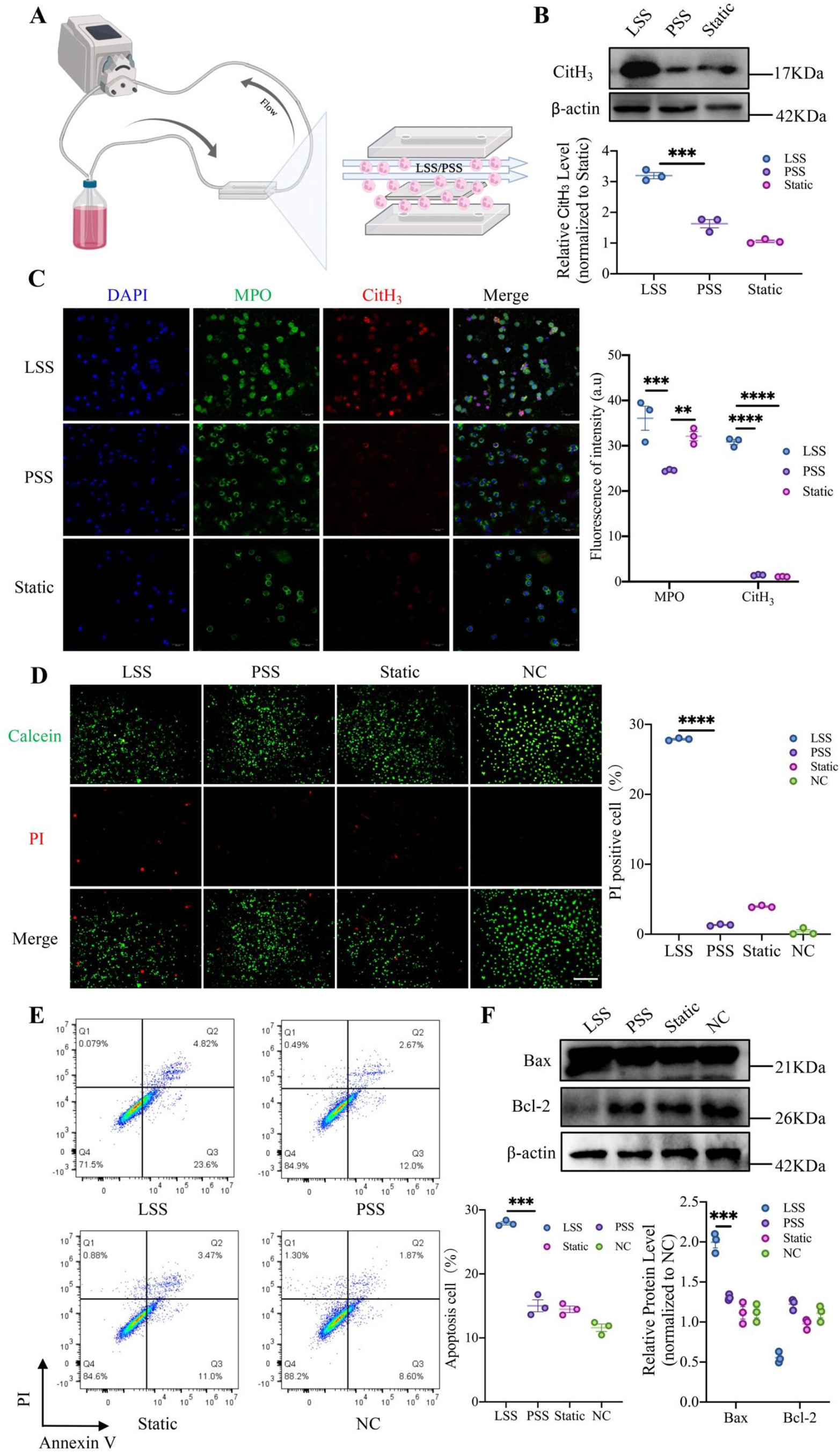
LSS promotes neutrophil extracellular traps (NETs) generation directly and aggravates endothelial cell apoptosis. (A) Schematic of shear stress treatment *via* parallel-plate flow chamber. (B and C) dHL-60 cells were treated with different shear stress (LSS, 5 dyne/cm^2^; PSS, 12 dyne/cm^2^; Static, 0 dyne/cm^2^) for 24 h. The expression levels of CitH_3_ were markedly upregulated under LSS. Scale bar, 50 μm. (D) Calcein (green) and PI (red) co-staining were detected in HUVECs which were co-cultured with different shear stress-induced dHL-60 cells for 24 h and quantified as total red fluorescent staining per visual field. Scale bar, 100 μm. (E) Flow cytometry was used to detect the apoptosis of HUVECs after co-cultured with different shear stress-induced dHL-60 cells for 24 h. The apoptosis rate was calculated *via* the positive rate of Annexin V-PI staining. (F) Bax and Bcl-2 protein expression were determined by Western blot. Band densities were quantitatively analyzed and normalized by β-actin. The data are presented as the mean ± SEM. (n=3). \*\**P* < 0.01, \*\*\**P* < 0.001, \*\*\*\**P* < 0.0001.

To further validate the effects of shear stress-induced dHL-60 cells on HUVECs, we co-cultured different shear stress-induced dHL-60 cells with HUVECs. As shown in Figure. 3D, LSS-induced dHL-60 cells promote more cell death of HUVECs. We further detected HUVEC apoptosis *via* flow cytometry, and found that LSS-induced dHL-60 cells exacerbated HUVEC apoptosis (Figure. 3E). After that, we compared apoptosis regulator Bcl-2 and Bax protein expression in different groups. LSS-induced dHL-60 cells downregulated Bcl-2 expression and upregulated Bax expression in HUVECs (Figure. 3F). Collectively, these findings suggest that LSS-induced dHL-60 cells have potent effects on regulating HUVECs apoptosis *in vitro*.

To better observe the interaction between neutrophils and endothelial cells under LSS, we seeded HUVECs on slides, which were assembled within the parallel-plate flow chamber system. dHL-60 cells flow through the system with the help of a pump. Immunofluorescence staining results show that LSS-induced dHL-60 cells were adhere to endothelial cells better than others (Figure. S3A). Meanwhile, the apoptosis rate of flowing cells in the LSS group was lower than that in the control group, which was consistent with the adhesion results (Figure. S3B).

### 3.4. LSS downregulates Piezo1 expression and elevated ROS

To further clarify the specific mechanism of neutrophils sense under shear stress, we first compared the mRNA and protein expression levels of Piezo1 in dHL-60 cells at LSS, PSS, and static respectively. Based on qPCR experiments, expression of Piezo1 was significantly decreased in dHL-60 cells subjected to LSS, compared with PSS and static. These results were also confirmed by Western blot analysis (Figure. 4A and 4B). Immunofluorescence showed that Piezo1 was dramatically down-regulated after being treated by LSS (Figure. 4C). Meanwhile, a significant increase in HDAC2 was found in agreement with decreased Piezo1 expression among the three groups (Figure. 4A and 4B). In addition, the intracellular Ca^2+^ concentration of dHL-60 cells was obviously decreased in LSS compared to PSS and statics (Figure. 4D). Interestingly, the intracellular ROS concentration also increased in LSS condition (Figure. 4E). These results imply that LSS may directly promote NETs generation through the Piezo1-HDAC2 axis.

**Figure 4.**
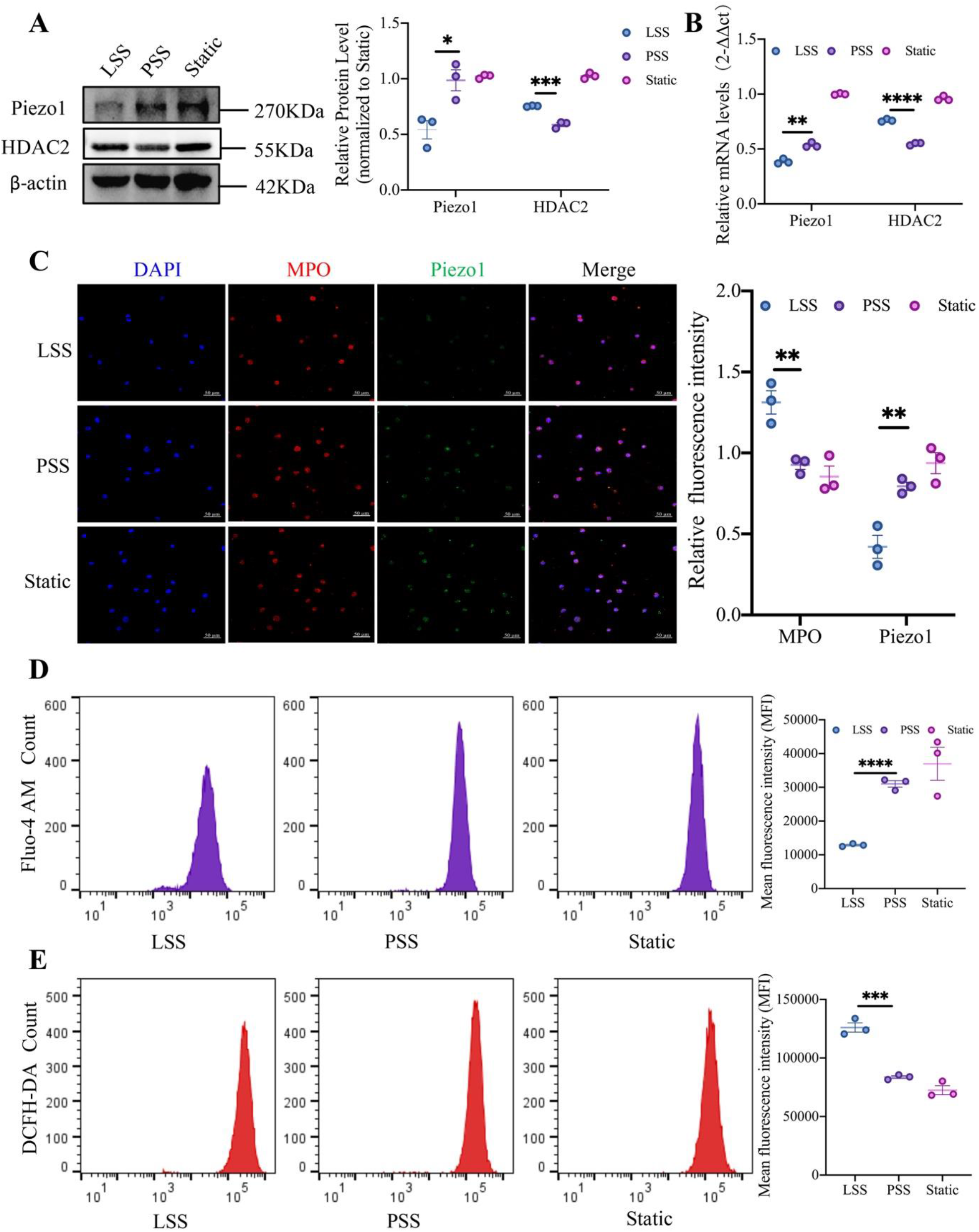
LSS downregulates Piezo1 and elevates reactive oxygen species (ROS) levels. dHL-60 cells were exposed to LSS or PSS for 24h. (A) Piezo1 and HDAC2 protein expression of dHL-60 cells were determined by Western blot. (B) Piezo1 and HDAC2 mRNA level of dHL-60 cells was determined by RT-qPCR, normalized to the amount of β-actin. (C) Immunofluorescent staining of the Piezo1 expression of dHL-60 cells. Scale bar, 50 μm. (D) Intracellular Ca^2+^ concentration of dHL-60 cells. (E) Intracellular ROS levels of dHL-60 cells. The data are presented as the mean ± SEM. (n=3). \**P* < 0.05, \*\**P* < 0.01, \*\*\**P* < 0.001.

### 3.5 Piezo1 knockdown promotes NETs generation in dHL-60 cells

To investigate whether the promotive effect of LSS on the NETs generation of dHL-60 cells was mediated by Piezo1, we knockdown Piezo1 in dHL-60 cells *via* siRNA transfection. Scramble siRNA served as control. A 50-75% knockdown efficiencies were verified by Western blot and immunofluorescence analysis prior to experiments (Fig. S4A and S4D). immunoblot demonstrated that the protein abundance of CitH_3_ was dramatically increased in the Piezo1-knockdown dHL-60 cells (Figure. 5A). Immunofluorescent staining with anti-CitH_3_ antibody showed that the numbers of CitH_3_^+^ cells were significantly increased in Piezo1-knockdown dHL-60 cells (Figure. 5B). In addition, a significant increase in HDAC2 was found in the Piezo1-knockdown dHL-60 cells both in protein and mRNA levels (Figure. 5A and 5C). As expected, Piezo1 knockdown also inhibited intracellular calcium influx and resulted in an increase in intracellular ROS levels (Figure. 5F and 5G).

**Figure 5.**
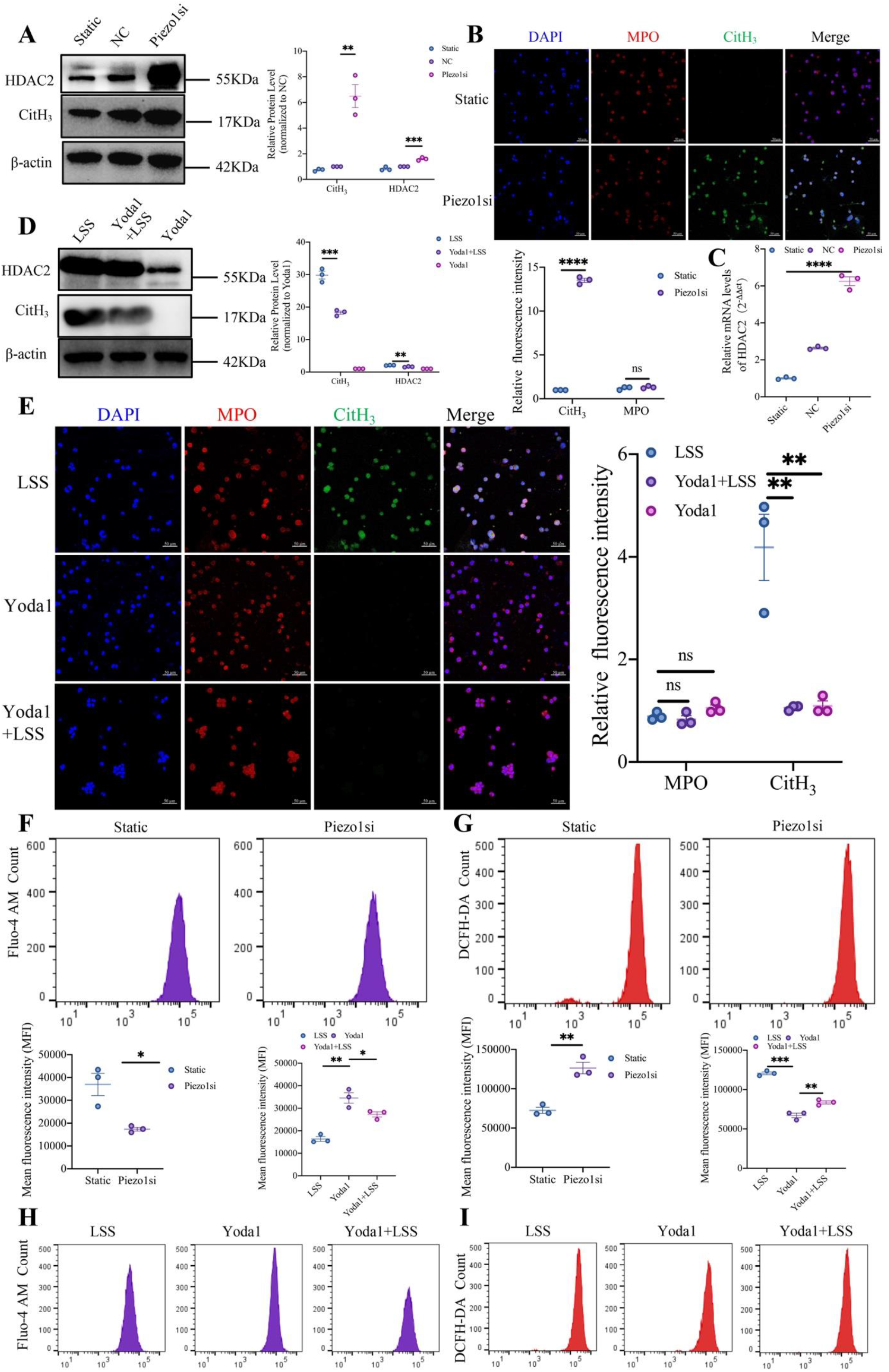
Piezo1 knockdown promotes neutrophil extracellular traps (NETs) generation in dHL-60 cells and Piezo1 overexpression attenuates LSS-induced dHL-60 cells NETosis *in vitro*. (A) Representative immunoblots and corresponding densitometry analysis of CitH_3_ and HDAC2 protein abundance in dHL-60 cells transfected with Piezo1 siRNA. (B) Immunofluorescence of CitH_3_ and MPO in dHL-60 cells transfected with Piezo1 siRNA. Scale bar: 50 μm. (C) HDAC2 mRNA level of Piezo1 knockdown dHL-60 cells was determined by RT-qPCR, normalized to the amount of β-actin. (D) Representative immunoblots and corresponding densitometry analysis of CitH_3_ and HDAC2 protein abundance in dHL-60 cells of LSS, Yoda1 and Yoda1+LSS group. (E) Immunofluorescence of CitH_3_ (green), MPO (red) and DPAI (blue) in dHL-60 cells of LSS, Yoda1 and Yoda1+LSS group. Scale bar: 50 μm. Flow cytometry analysis of intracellular Ca2+ (F) and ROS concentrations (G) of dHL-60 cells transfected with Piezo1 siRNA. Flow cytometry analysis of intracellular Ca2+ (H) and ROS concentrations (I) of dHL-60 cells treated with Yoda1. The data are presented as the mean ± SEM. (n=3). \**P* < 0.05, \*\**P* < 0.01, \*\*\**P* < 0.001, \*\*\*\**P* < 0.0001.

### 3.6 Piezo1 overexpression attenuates LSS-induced dHL-60 cell NETosis in vitro

To further demonstrate Piezo1’s role in regulating NETs formation in dHL-60 cells, Yoda1, the agonist of Piezo1, was added to dHL-60 cells subjected to LSS or not. Piezo1 protein levels were measured by western blot and Immunofluorescence analysis (Figure. S4B and S4C). Piezo1 mRNA levels were measured by RT-qPCR (Figure. S4E). Interestingly, although Yoda1 (1 μM, 24h) did not alter NETs generation in static dHL-60 cells, it can significantly reverse LSS-induced NETs formation (Figure. 5D). This result was further confirmed by the immunofluorescence staining. (Figure. 5E). Meanwhile, a significant decrease in HDAC2 was found in agreement with increased Piezo1 expression among the three groups (Figure. 5D). Moreover, treatment of a Piezo1 agonist Yoda1 increased calcium influx and decreased ROS levels in dHL-60 cells (Figure. 5H and 5I). These results further demonstrate that Piezo1 play an essential role in sensing mechanical loading in neutrophils.

### 3.7 HDAC2 inhibition affects little of the expression of piezo1 and NETosis

To further verify the relationship between Piezo1 and HDAC2, 5 mM VPA, a specific HDAC2 inhibitor, was added to dHL-60 cells for 6 hours. HDAC2’s expression levels were identified by RT-qPCR, western blot and immunofluorescence analysis (Figure. S5). VPA alone had no effect on NETosis in dHL-60 cells. However, VPA could reverse piezo1-knockdown-induced NETosis (Figure. 6A). This result was further confirmed by the immunofluorescence staining. (Figure. 6C). VPA had little effect on the expression of Piezo1(Figure. 6A and 6D) and intracellular Ca^2+^ concentration (Figure. 6G), suggesting that piezo1 and Ca^2+^ may be upstream regulators of HDAC2. VPA reversed LSS-induced elevation of ROS levels (Figure. 6H), suggesting that HDAC2 promoted NETosis probably through ROS.

**Figure 6.**
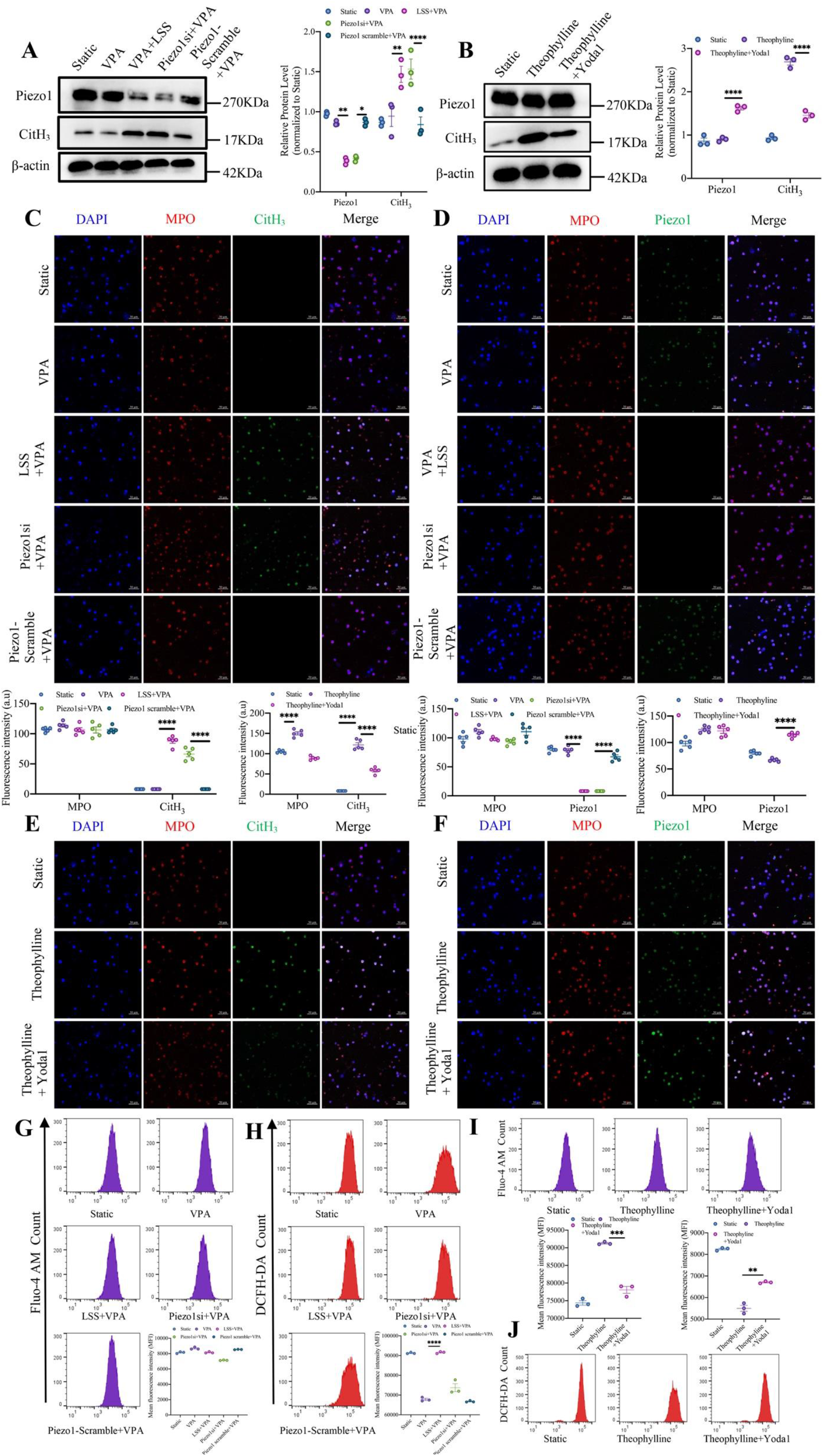
HDAC2 inhibition affects little of the expression of piezo1 and NETosis. Static and Piezo1 knockdown dHL-60 cells treated with valproic acid sodium (VPA) or not. Yoda1 attenuates NETosis induced by HDAC2 overexpression in dHL-60 cells. Static dHL-60 cells treated with Theophylline or not. (A and B) Representative immunoblots and corresponding densitometry analysis of Piezo1 and CitH_3_ protein abundance in dHL-60 cells under different treatments. Immunofluorescence of CitH_3_ (green) (C and E), Piezo1 (green) (D and F), MPO (red), and DPAI (blue) in dHL-60 cells under different treatments. Scale bar: 50 μm. Flow cytometry was used to detect the intracellular Ca2+ (G and I) and ROS (H and J) concentrations of dHL-60 cells under different treatments. The data are presented as the mean ± SEM. (n=3). \**P* < 0.05, \*\**P* < 0.01, \*\*\**P* < 0.001.

### 3.8 HDAC2 overexpression affects intracellular Ca^2+^ and NETosis

Further, we investigated the effect of overexpression of HDAC2 on NETosis. Theophylline (5μM, 6h), an agonist of HDAC2, was used to up-regulate the expression of HDAC2 in dHL-60 cells. HDAC2 mRNA was measured by RT-qPCR and protein levels were measured by western blot and Immunofluorescence analysis (Figure. S5). In static cultivation of dHL-60 cells, Theophylline could strongly induce the formation of NETs. Meanwhile, Yoda1treatment attenuated the promoting effect of Theophylline on NETosis (Figure. 6B). This was consistent with the immunofluorescence results (Figure. 6E). The increased HDAC2 did not significantly affect the expression of piezo1 (Figure. 6B and 6F) but obviously reduced the intracellular Ca^2+^ concentration (Figure. 6I). It suggests that piezo1 is indeed an upstream regulator of HDAC2 and there may be an interaction between calcium ions and HDAC2. ROS levels were significantly increased upon Theophylline stimulation, which could also be reduced by Yoda1 (Figure. 6J). The result reveals that ROS is an effector of HDAC2 and is a key molecule that induces NETs generation. In aggregate, these data indicate that LSS triggers NETs formation may through Piezo1, which downregulates intracellular Ca^2+^ levels, upregulate HDAC2 expression and facilitate ROS generation.

## 4. Discussion

Atherosclerosis is a complex pathologic process involved in interplay of lipid metabolism alterations, endothelial dysfunction, inflammation, oxidative stress, as well as hemodynamic factors^20, 21^. Recent research has highlighted the critical role of lipid-driven chronic inflammation in atherogenesis^22^, especially the early involvement of neutrophils and its distinctive death process, NETosis^23^. However, most of the studies focused on the effect of shear stress on endothelial cells or smooth muscle cells and less attention has been placed on how it induces NETs formation. In this study, we demonstrated that LSS directly promotes NETosis through the piezo1-HDAC2 axis in atherosclerosis progression.

Although NETs were reported as a new defense mechanism of neutrophils in face of pathogen^24^, most researches favor that NETs promote the development of atherosclerosis. NETs were found in human and mouse carotid plaques and was reported to be associated with increased atherosclerosis progression^7, 25^. NETs impede atherosclerosis resolution by increasing plaque inflammation and propagating endothelial dysfunction^26^. The increased plaque stability and decreased plaque necrotic area upon DNase1-treatment also indicated the toxic effects of NETs in atherosclerotic mice^27^. In our study, we observed a higher aggregation degree of NETs in the plaques with longer fat feeding in ApoE^−/−^ mice. Daily intraperitoneal injection with GSK484, a PAD4 inhibitor, could significantly reduce the formation of plaque and the production of NETs in experimental animal models. However, under the same high-fat diet, the reduction of NETs did not cause appreciable effects on overall circulating lipid levels, indicating that NETs are involved in lipid metabolism especially the deposition of lipids in the vascular wall, rather than lipid production. We speculate that this is associated with the scaffold function of NETs^28^. Thus, the formation of NETs may be the initial factor for the deposition of fat particles in the vascular wall.

However, the trigger of NETosis during atherosclerosis is still unclear. Various stimuli have been reported to induce NETs generation, which could be broadly divided into three major classes including pathogenic microorganisms, inflammatory factors, and other small particles^29^. As far as we know, there is no report about the correlation between shear stress and NETosis. In this study, we employed a partial carotid ligation mouse model that rapidly induces gene expression changes, endothelial inflammation, and atherosclerosis^30^. We observed significantly increased generation of NETs in carotid plaques under LSS. Furthermore, we found that NETs could be directly induced by LSS using *in vitro* parallel plate flow chamber experiments. It is surprising that the magnitude of flow shear stress affects the activation state of neutrophils. It also suggests that there may be molecules on neutrophils that directly sense shear stress instead of conventional molecules which converts drag forces of flowing blood into bond tension.

Several molecules have been identified to play roles in neutrophil mechanotransduction, including cell adhesion molecules (*eg:* integrins and cadherins), transmembrane receptors (*eg:* G-protein coupled receptors)^31^, cytoskeletal (*eg:* F-actin)^32^, and mechanosensitive ion channels such as TRPC6, TRPM7, TRPV family^33^. Piezo1 channel, a newly discovered mechanically sensitive ion channel (MSC), has been reported as an effective mechanical sensor for cells, such as cortical astrocytes^34^, cardiomyocytes^35^, alveolar type I epithelial cells^36^. Our study demonstrates Piezo1 may be a novel shear stress sensor for neutrophils, and play a crucial role in LSS-induced NETosis. dHL-60 cells under LSS showed lower Piezo1 expression and enhanced formation of NETs, which could be alleviated by Yoda1-induced Piezo1 overexpression. Meanwhile, Piezo1 knockdown could also promot NETs generation in static cells. These results support that Piezo1 is crucial for maintaining the physiological homeostasis of neutrophils, and is essential for NETosis. Our study enriched the mechanosensor of ion channels on neutrophils.

Piezo1 channel is a kind of cation-selective mechanical channel with a selectivity sequence of Ca^2+^ > K^+^ > Na^+^ > Mg^2+ 37^. Intracellular Ca^2+^ imbalance is known to directly induce NETs formation by activating the PAD4 enzyme, driving histone citrullination and chromatin decondensation through Nox2 independent mechanism^38^. Therefore, we investigated the change of intracellular Ca^2+^ concentration under LSS. We found that LSS-induced downexpression of the Piezo1 caused reduction of intracellular Ca^2+^ concentration, while the agonist effects of Yoda1 could significantly elevate the intracellular Ca^2+^ concentration inhibited by LSS. The results illustrate the role of intracellular Ca^2+^ concentration in LSS-induced NETs generation.

HDACs belong to two families, based on the dependency on zinc or nicotinamide adenine dinucleotide (NAD+) for their activities^39^. It has been reported that HDACs play critical roles in driving NETs formation in human and mouse neutrophils. Inhibition of HDACs in mice protects against microbial-induced pneumonia and septic shock^40^. In our reserch, Knockdown of Piezo1 can significantly enhance the expression of HDAC2, while changing the expression of HDAC2 has little effect on Piezo1. However, the detailed mechanism of how Piezo1 regulates HDAC2 and whether Ca^2+^ participate in HDAC2 regulation is not clear, and requires further exploration. NETosis was previously reported to be of two types: ROS dependent and ROS independent^41^. Our study shows that LSS may induce ROS-dependent NETosis *via* HDAC2, which could promote the accumulation of total cellular ROS^42^.

According to previous studies, the mechanism of NETs participates in atherosclerosis is probably by directly mediating organ and endothelial toxicity^7, 26, 43^. NETs-induced endothelial stress and dysfunction are dependent on TLR2 activation and the release of interferon-*α*(IFN-*α*)^44^, or CitH3^45^. Similarly, damaged endothelial cells upregulate different chemokines (CCL5, CXCL1, CCL2) and adhesion molecules (E-selectin, P-selectin, ICAM-1, VCAM-1) which enhance the adhesion and infiltration of neutrophil into the intimal space^46, 47^. We further examined the effect of shear stress-induced dHL-60 cells on endothelial cells through a series of coculture experiments. In agreement with previous studies, we observed decreased activity, increased apoptosis, and stronger adhesion to NETs of endothelial cells when exposed to LSS-induced dHL-60 cells.

In summary, our study shows that LSS-induced NETs generation may play an crucial role in the development of atherosclerosis. In animal experiments, NETs inhibition significantly reduced plaque formation and impeded the progression of atherosclerosis. In cell experiments, LSS promotes NETosis by downregulate piezo1 expression to imbalance intraceullar Ca^2+^ concentration and increase HDAC2 expression. This intriguing new concept may enlighten new mechanisms involved in plaque erosion and might facilitate the development of new treatments for atherosclerotic diseases, especially vascular bifurcation lesions. It could also help to further understand the mechanism of NETosis and NETosis-related inflammation.

## Nonstandard Abbreviations and Acronyms

NETs: neutrophil extracellular traps
HFD: high-fat diet
LSS: lower shear stress
PSS: physiological shear stress
HDAC2: histone deacetylase 2
ROS: reactive oxygen species
PAD4: peptidyl arginine deiminase-4
dHL-60: differentiated human promyelocytic leukemia HL-60
LCA: left carotid artery
RCA: right carotid artery
HUVECs: human umbilical vein endothelial cells
CitH_3_: citrullination of histone H3
MPO: myeloperoxidase
i.p.: intraperitoneally
VPA: valproic acid sodium

## Acknowledgments

We thank the staff of the Center of Smart Laboratory and Molecular Medicine of Chongqing University and the Department of Cardiovascular Surgery, Research Institute of Surgery, Daping Hospital, Army Medical University. We especially appreciate Dr. Yanyun Wang and Prof. Juhui Qiu of the School of BioEngineering, Chongqing University for their support and suggestions on this study.

## Supplemental Material

Supplemental Methods

Tables S1–S2

Figure S1-S5

